# Mps2 links Csm4 and Mps3 to form a telomere-associated LINC complex in budding yeast

**DOI:** 10.1101/2020.05.30.125559

**Authors:** Jinbo Fan, Hui Jin, Bailey A. Koch, Hong-Guo Yu

## Abstract

The linker of the nucleoskeleton and cytoskeleton (LINC) protein complex is composed of a pair of transmembrane proteins: the KASH-domain protein localized to the outer nuclear membrane and the SUN-domain protein to the inner nuclear membrane. In budding yeast, the sole SUN-domain protein, Mps3, is thought to pair with either Csm4 or Mps2, two KASH-like proteins, to form two separate LINC complexes. Here we show that Mps2 mediates the interaction between Csm4 and Mps3 to form a heterotrimeric telomere-associated LINC (t-LINC) in budding yeast meiosis. Mps2 binds to Csm4 and Mps3, and all three are localized to the telomere. Telomeric localization of Csm4 depends on both Mps2 and Mps3; in contrast, Mps2’s localization depends on Mps3 but not Csm4. Mps2-mediated t-LINC regulates telomere movement and meiotic recombination. By ectopically expressing *CSM4* in vegetative yeast cells, we reconstitute the heterotrimeric t-LINC and demonstrate its ability to tether telomeres. Our findings therefore reveal the heterotrimeric composition of t-LINC in budding yeast and have implications for understanding LINC variant formation.

## Introduction

The linker of the nucleoskeleton and cytoskeleton (LINC) protein complex has emerged as a key regulator for a diverse range of nuclear activities that include chromosome movement, nuclear positioning, and gene expression (Tapley and Starr, 2013; Chang et al., 2015; Burke, 2018). The canonical LINC complex is composed of a pair of transmembrane proteins, the SUN (**S**ad1 and **UN** C-84) protein localized to the inner nuclear membrane (INM) and the KASH (**K**larsicht, **A**NC-1 and **S**yne/Nesprin **h**omology) protein localized to the outer nuclear membrane (ONM)(Hagan and Yanagida, 1995; Welte et al., 1998; Malone et al., 1999; Starr and Han, 2002). With SUN-KASH interaction taking place in the perinuclear space, the LINC complex not only bridges the INM and ONM, but also connects the cytoskeleton to the nucleoplasm for allowing transduction of mechanical forces through the nuclear envelope (Starr and Fridolfsson, 2010). At least five SUN-domain proteins and six KASH-domain proteins have been found in mammals (Rajgor and Shanahan, 2013; Duong et al., 2014; Chang et al., 2015; Nishioka et al., 2016), potentially forming a diverse number of LINC complex variants. Similarly, SUN- and KASH-like proteins are prevalent in land plants (Zhou and Meier, 2013; Gumber et al., 2019). LINC proteins are believed to form heterodimeric hexamers and possibly higher-ordered protein arrays for force transmission and function (Luxton et al., 2010; Sosa et al., 2012; Wang et al., 2012; Chang et al., 2015), but the stoichiometry of how they are assembled *in vivo* remains to be further determined.

In budding yeast, Mps3 is the sole SUN-domain protein, which is concentrated at the yeast centrosome (Jaspersen et al., 2002), often called the spindle pole body (SPB). Mps3 also localizes to the INM (Jaspersen et al., 2002). Budding yeast lacks a canonical KASH-domain protein, but possesses two ONM-localized KASH-like proteins: Mps2, present in both mitosis and meiosis (Winey et al., 1991), and Csm4, a meiosis-specific protein (Conrad et al., 2008; Kosaka et al., 2008; Koszul et al., 2008; Wanat et al., 2008). The genes encoding Mps2 and Csm4 are considered paralogs, but they differ drastically in protein size and show limited similarity at the amino acid level (Fig 1A). The current notion posits that in mitosis and presumably also in meiosis, Mps3 pairs with Mps2 at the centrosome to form a centrosome-associated LINC (Jaspersen et al., 2006; Chen et al., 2019), to which we refer as c-LINC. In meiosis, Mps3 pairs with Csm4 to form a telomere-associated LINC (t-LINC) (Conrad et al., 2008; Kosaka et al., 2008; Koszul et al., 2008; Wanat et al., 2008).

**Figure 1.**
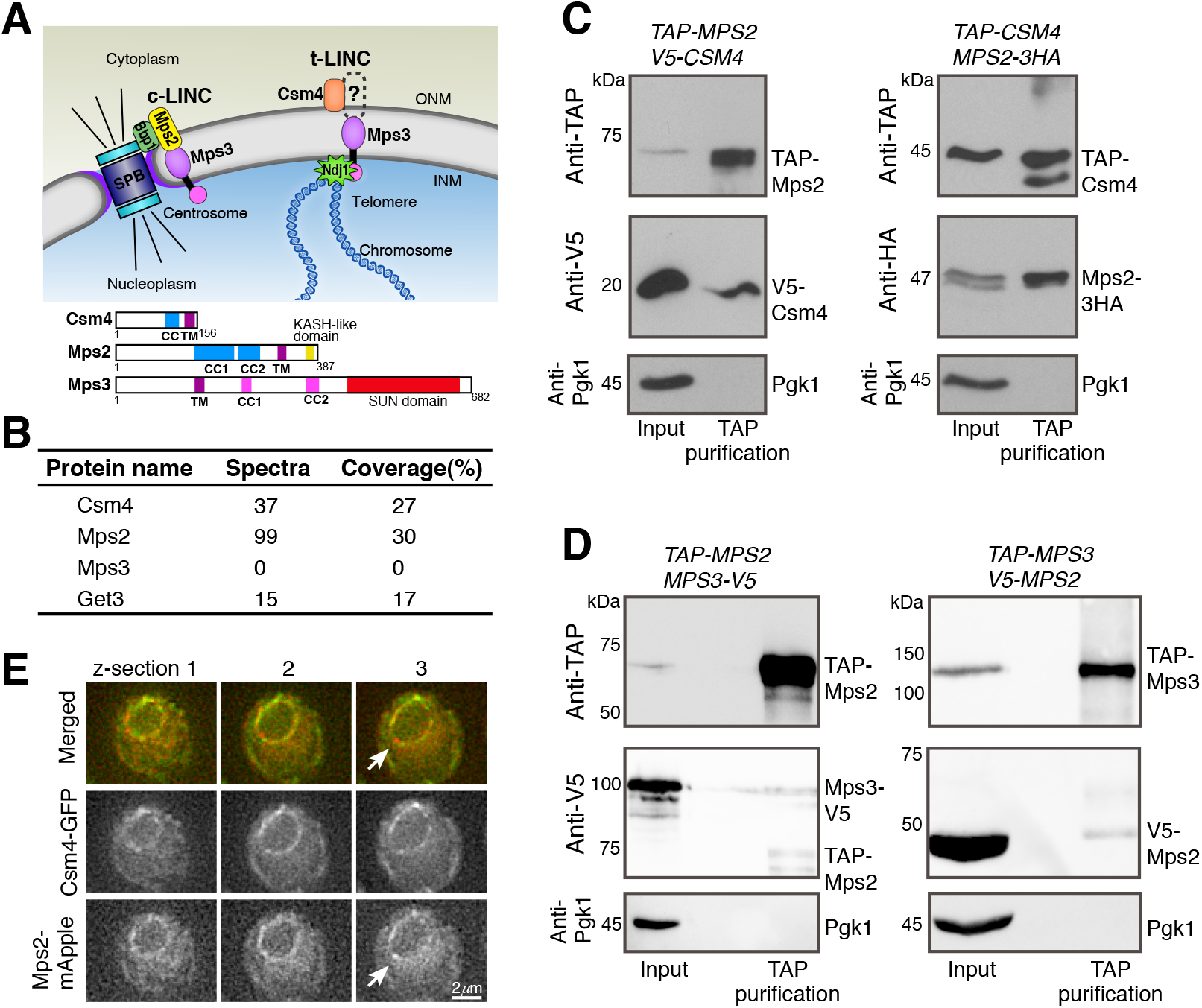
Meiotic Mps2 binds to Csm4 and Mps3. (**A**) Schematic diagram showing the composition and location of c-LINC and t-LINC in budding yeast. Protein structures of Csm4, Mps2 and Mps3 are shown at the bottom. INM, inner nuclear membrane; ONM, outer nuclear membrane; CC, coiled coil; TM, transmembrane domain. (**B**) List of representative proteins copurified with TAP-Csm4. (**C**) Reciprocal immunoprecipitation showing Mps2-Csm4 interaction. The level of Pgk1 serves as a control for affinity purification. (**D**) Reciprocal immunoprecipitation showing Mps2-Mps3 interaction. Note that the anti-V5 antibody also recognizes TAP-Mps2. (**E**) Localization of Mps2 and Csm4 at prophase I. Three continuous optical sections are shown. Arrows point to the putative localization of Mps2 to the spindle pole body (SPB). Note that both Csm4 and Mps2 localize to the nuclear periphery. Red, Mps2-mApple; green, Csm4-GFP.

Classified as single-pass type II transmembrane proteins, Mps2 and Mps3 most likely interact in the perinuclear space through their corresponding C-terminal KASH-like and SUN domains, about 60 and 190 amino acids in size, respectively (Fig 1A). Like Mps3, Mps2 is also concentrated at the SPB, where it binds to additional SPB components to form a subcomplex that regulates SPB insertion into the nuclear envelope (Munoz-Centeno et al., 1999; Schramm et al., 2000). In vegetative yeast cells, Mps3 plays additional roles in DNA double-strand break repair by anchoring chromosomes to the nuclear periphery (Kalocsay et al., 2009; Oza et al., 2009). Whether Mps2 has a similar role in recombination and chromosome tethering remains unclear.

During meiosis, Mps3 is required for tethering telomeres to the nuclear envelope in addition to its roles in SPB duplication and separation (Conrad et al., 2007; Conrad et al., 2008; Li et al., 2015; Li et al., 2017). At meiotic prophase I, the N-terminus of Mps3, which is located in the nucleoplasm, binds to Ndj1, a telomere-associated protein (Chua and Roeder, 1997; Conrad et al., 1997), whereas its C-terminal SUN domain, located in the perinuclear space, has been proposed to bind to Csm4 (Fig 1A and Conrad et al., 2008). Therefore, the t-LINC, composed of Csm4 and Mps3 at a minimum, is capable of linking telomeres to the cytoplasmic actin filaments, which provide the mechanical forces necessary for meiotic telomere movement (Koszul et al., 2008). This t-LINC-dependent motility mediates the configuration of the telomere bouquet and can drastically deform the nucleus at prophase I (Conrad et al., 2008; Koszul et al., 2008). However, Csm4 is a tail-anchored membrane protein with its predicted transmembrane domain located at the very end of the C-terminus, exposing merely a three-amino acid tail in the perinuclear space (Fig 1A). For canonical KASH proteins, their tails are usually ~30 amino acids long (Sosa et al., 2012; Wang et al., 2012). In the absence of a KASH-like domain as found in Mps2, how Csm4 interacts with Mps3 in the perinuclear space remains unclear.

We hypothesize that another unidentified factor is required to mediate Csm4 and Mps3 interaction to form the t-LINC. We report here that Mps2 mediates the interaction between Csm4 and Mps3 to form a heterotrimeric t-LINC that tethers telomeres and regulates nuclear dynamics in budding yeast meiosis. Using a combined cytological and genetic approach, we show that Mps2 is a major binding partner of Csm4 and a telomere-associated protein. Furthermore, by ectopically expressing *CSM4* in vegetative yeast cells, we have reconstituted the heterotrimeric t-LINC and demonstrated its ability to tether telomeres. Our findings therefore reveal the heterotrimeric composition of the yeast t-LINC.

## Results and Discussion

### Meiotic Mps2 is a major binding partner of Csm4

To test our hypothesis that an unidentified factor mediates the interaction between Csm4 and Mps3, we performed TAP-Csm4 protein affinity purification, followed by mass-spectrometry-based protein identification (Fig 1B). Purification of Csm4 was confirmed by 37 identified peptide spectra that belonged to Csm4 and covered 27% of its amino acid sequence (Fig 1B). As expected, Get3, which is required for insertion of tail-anchored proteins into the ER membrane (Schuldiner et al., 2008), was copurified with TAP-Csm4 (Fig 1B). However, Mps3 was not identified (Fig 1B). Unexpectedly, a major protein copurified with TAP-Csm4 was Mps2, which showed 99 peptide spectra that covered 30% of the Mps2 protein sequence (Fig 1B). This and additional findings described below prompted us to propose that Mps2 is a t-LINC component in budding yeast.

To further determine the interaction between t-LINC components, we performed reciprocal immunoprecipitation, followed by western blotting (Fig 1C and 1D). Yeast cells were induced to undergo synchronous meiosis, and arrested at prophase I by way of *ndt80Δ* (Xu et al., 1995). V5-Csm4 was copurified with TAP-Mps2; reciprocally, Mps2-3HA was purified by immunoprecipitation of TAP-Csm4 (Fig 1C). By fluorescence microscopy, we confirmed that Mps2 and Csm4 colocalized to the nuclear periphery during meiosis (Fig 1E). In addition, Mps3-V5 was copurified with TAP-Mps2; V5-Mps2 was copurified with TAP-Mps3 (Fig 1D). With this TAP method, however, we did not observe a direct Csm4-Mps3 interaction (Fig 1B and unpublished data). In summary, we have revealed the interaction between Mps2 and Csm4 and confirmed the interaction between meiotic Mps2 and Mps3, which has not been reported previously.

### Mps2 is required for meiotic cell progression

To better understand the role of Mps2 in meiosis, first, we determined Mps2 localization by time-lapse fluorescence microscopy (Fig 2A). As expected, meiotic Mps2 was found at the SPB, evidenced by its colocalization with the SPB marker Tub4, which first appeared as a focus, then separated from one focus to two foci in meiosis I and two to four in meiosis II (Fig 2A and Supplemental Fig 1A). In early meiosis II, Mps2 was preferentially associated with the newly duplicated SPB, which displayed a weaker Tub4-mApple signal (Fig 2A, t=70) due to a slower fluorescence maturation time of mApple than that of GFP. Importantly, Mps2 localized around the nuclear periphery (Fig 1E, 2A and Supplemental Fig 2A). Before SPB separation in meiosis I, distribution of Mps2 at the nuclear envelope appeared uneven, at times displaying high occupancy to only half or less than half of the nuclear periphery (Fig 2A, t=−30 min for an example and Supplemental Fig 2A). This polarized localization of Mps2 to a certain region of the nuclear envelope is similar to that of Csm4 at prophase I (Kosaka et al., 2008; Wanat et al., 2008) and lends support to the idea that like Csm4, Mps2 is a component of the t-LINC.

**Figure 2.**
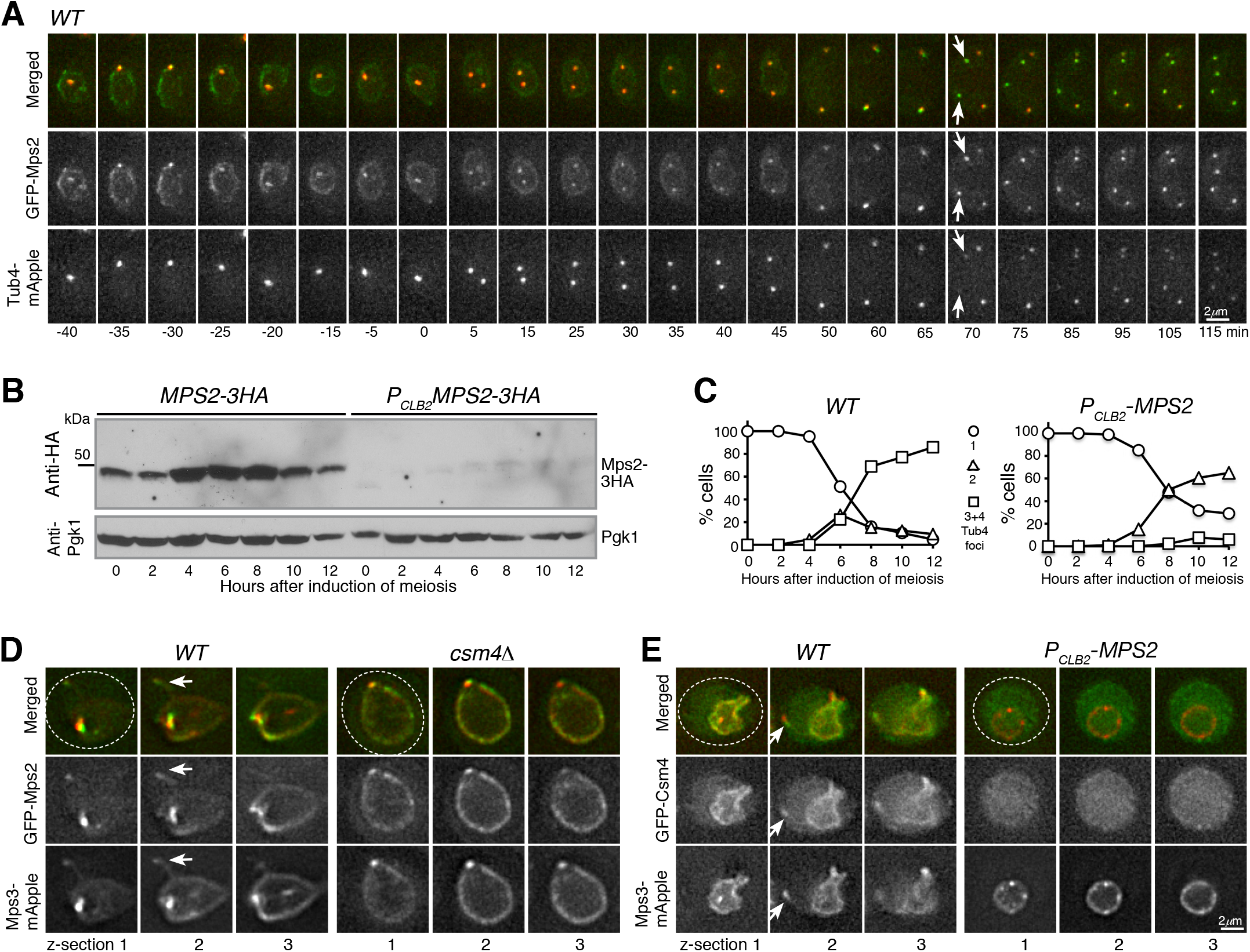
Mps2 is required for meiotic cell progression and regulates Csm4 localization. (**A**) Time-lapse fluorescence microscopy showing GFP-Mps2 localization during meiosis. Tub4-mApper serves as a marker for the SPB. Projected images of 12 z-sections are shown. Arrows point to the newly duplicated SPBs in meiosis II. Time in minutes is shown at the bottom. Time zero refers to the onset of SPB separation in meiosis I. Note the uneven localization of Mps2 to the nuclear periphery in meiosis I. Red, Tub4-mApple; green, GFP-Mps2. (**B**) Protein level of Mps2 in budding yeast meiosis. Yeast cells were induced to undergo synchronous meiosis; cell aliquots were withdrawn at indicated times. The level of Mps2-3HA was probed by an anti-HA antibody. The level of Pgk1 serves as a loading control. Note that Mps2 was largely depleted in *P_CLB2_*-*MPS2* cells. (**C**) SPB separation in wild-type (*WT*) and *P_CLB2_*-*MPS2* cells during meiosis. Tub4-mApple serves as a marker for the SPB. At least 100 cells were counted at each time point. Three biological replicates were performed, shown is a representative. Note that *P_CLB2_*-*MPS2* cells were stopped with only two SPBs. (**D**) Colocalization of Mps2 and Mps3. Note that GFP-Mps2 (green) and Mps3-mApple (red) remain bound to the nuclear periphery in the *csm4Δ* cell. (**E**) Mps2 is required for nuclear localization of Csm4. Note that the nucleus becomes a sphere in the *P_CLB2_*-*MPS2* cell. Three continuous optical sections are shown in D and E. Arrows point to the nuclear protrusion. Dashed lines show the overall cell shape. Red, Mps3-mApple; green, GFP-Csm4.

Next, we generated a meiosis-specific Mps2-depletion allele, *P_CLB2_*-*MPS2*, of which the endogenous *MPS2* promoter was replaced by that of *CLB2* (Fig 2B, and Lee and Amon, 2003). During meiosis, the level of Mps2 increased 4 hours after the induction of meiosis, which roughly corresponded to meiosis I. On the other hand, depletion of meiotic Mps2 appeared to be near completion in *P_CLB2_*-*MPS2* cells (Fig 2B). In the absence of Mps2, meiosis I occurred in more than 60% of the cells, as determined by the separation of Tub4-mApple from one focus to two foci (Fig 2C and Supplemental Fig 1A-E). More than 80% of wild-type cells formed four Tub4-mApple foci upon completion of meiosis; in contrast, less than 5% of *P_CLB2_*-*MPS2* cells displayed three or four Tub4-mApple foci (Fig 2C and Supplemental Fig 1B and 1D), indicating that Mps2 is required for meiotic cell progression. Abolishing meiotic recombination by way of *spo11-Y135F* (Keeney et al., 1997) did not rescue the SPB separation defect in *P_CLB2_*-*MPS2* cells (Supplemental Fig 1C and 1E). Furthermore, SPB separation appeared normal in *csm4Δ* cells (Supplemental Fig 1F and 1G). These findings indicate that meiotic Mps2 has additional functions outside of the t-LINC complex, and is likely involved in c-LINC-mediated meiotic SPB duplication as it is in mitosis, a topic for future study.

### Mps2 is required for nuclear localization of Csm4 but not for Mps3

To further test our hypothesis that Mps2 is a component of the t-LINC complex, we determined the interdependency of t-LINC components for their roles in nuclear envelope localization. We focused on meiotic cells at prophase I when t-LINC is active in regulating telomere bouquet formation and meiotic recombination (Conrad et al., 2008; Kosaka et al., 2008; Koszul et al., 2008; Wanat et al., 2008). By fluorescence microscopy, we observed that Mps2 and Mps3 colocalized to the nuclear periphery and to the SPB (Fig 2D). Similarly, their localization to the nuclear periphery appeared uneven at prophase I (Fig 2D). Because t-LINC mediates actin-based motility at the nuclear periphery (Koszul et al., 2008), the nuclear envelope formed membrane protrusions and the nuclear shape became irregular at prophase I (Fig 2D, 2E, and Conrad et al., 2008; Koszul et al., 2008). As expected, Mps2, Mps3 and Csm4 all were present at the leading edge of the nuclear protrusions (Fig 2D and 2E). In the absence of Csm4, the meiotic nucleus appeared as a sphere and lacked nuclear protrusions (Fig 2D and Conrad et al., 2008; Koszul et al., 2008). Mps2 and Mps3 remained bound to but were distributed evenly around the nuclear periphery in *csm4Δ* cells, demonstrating that association of both Mps2 and Mps3 with the nuclear envelope is independent of Csm4 but their polarized nuclear localization depends on Csm4. In Mps2-depleted cells, Csm4, but not Mps3, was no longer detectable at the nuclear periphery, and the meiotic nucleus became spherical without any visible membrane protrusions (Fig 2E). As in *csm4Δ* cells, Mps3 was distributed evenly around the nuclear periphery when Mps2 was absent (Fig 2E, right panels). Finally, we found that localization of Mps2 to the nuclear periphery, but not to the SPB, was impaired when Mps3 was depleted in yeast meiosis (Supplemental Fig 2A), indicating that Mps3 regulates the association of Mps2 with the nuclear envelope. Consequently, nuclear protrusions were not observed in cells depleted with meiotic Mps3 (Supplemental Fig 2B). Taken together, our findings demonstrate that localization of Csm4 to the nuclear periphery depends on Mps2, but not vice versa. Furthermore, Mps2 and Csm4 both are essential for generating nuclear protrusions, a major t-LINC activity, at prophase I.

### Meiotic Mps2 is a telomere-associated protein

If Mps2 is a component of the t-LINC, we reasoned that Mps2 would localize to the telomere, as Csm4 and Mps3 do (Conrad et al., 2007; Conrad et al., 2008). We therefore performed surface nuclear spreads, in which telomere-associated proteins can be determined by immunofluorescence (Fig 3). As a reference, cohesin Rec8 was used as a marker for the meiotic chromosome axis (Fig 3A-3D). We found that Mps2 colocalized with Ndj1 at the telomeres (Fig 3A). In addition, both Mps2 and Csm4 were colocalized at the chromosome ends (Fig 3B). Therefore, meiotic Mps2 is a telomere-associated protein.

**Figure 3.**
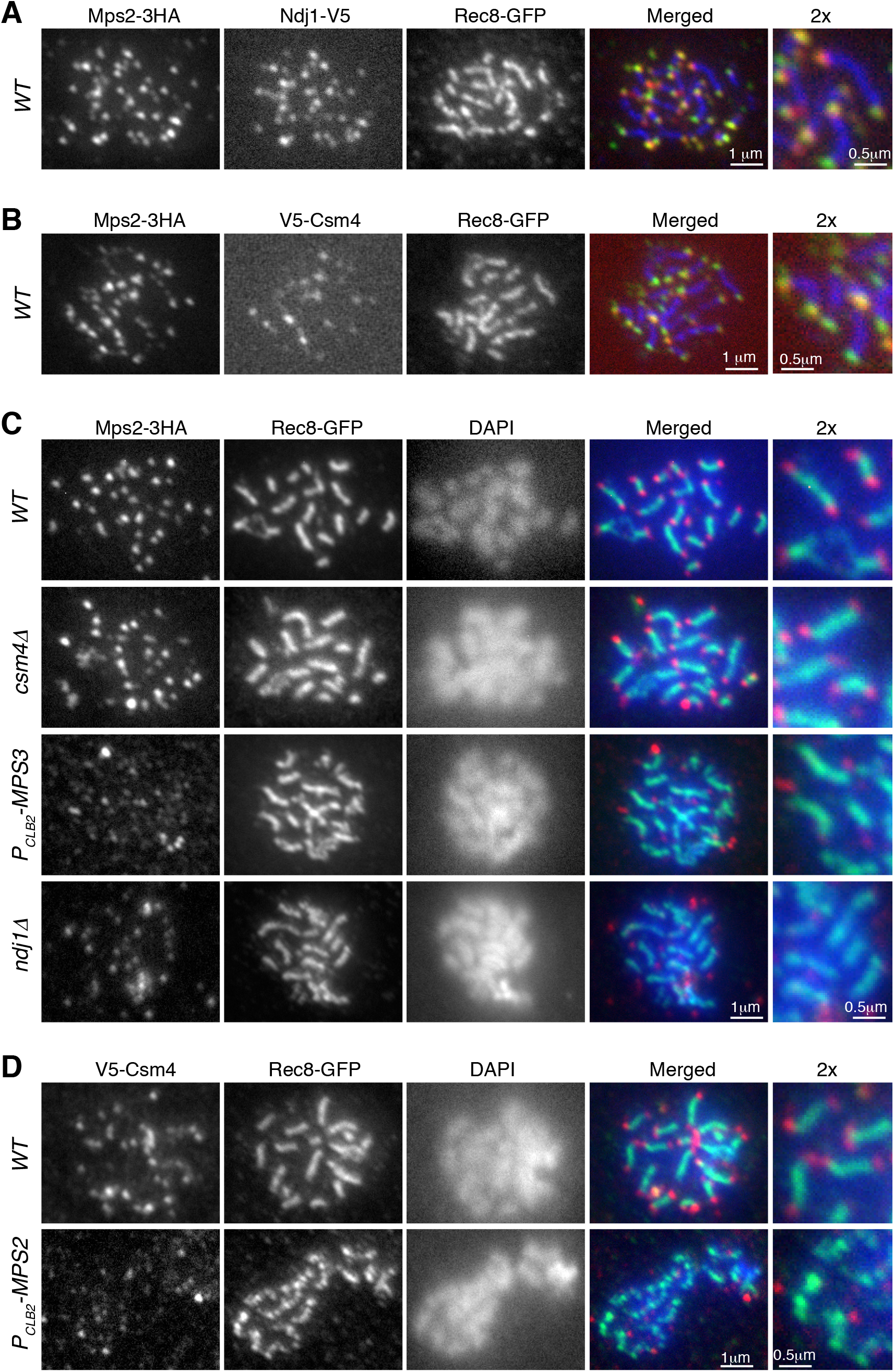
Mps2 is a telomere-associated protein. Meiotic cells were harvested for nuclear spreads, followed by immunofluorescence to probe V5, HA and GFP tagged proteins. DAPI stains DNA. Rec8 is used to mark the chromosome axis. Enlarged views (2x) are shown to the right. (**A**) Representative cell showing colocalization of Mps2 with Ndj1 to meiotic telomeres. Red, Ndj1-V5; green, Mps2-3HA; blue, Rec8-GFP. (**B**) Representative cell showing colocalization of Mps2 and Csm4 at telomeres. Red, V5-Csm4; green, Mps2-3HA; blue, Rec8-GFP. (**C**) Representative cells showing telomeric localization of Mps2 depends on Mps3 and Ndj1 but not Csm4. Red, Mps2-3HA; green, Rec8-GFP; blue, DAPI. (**D**) Representative cells showing telomeric localization of Csm4 depends on Mps2. Note that chromosome axes appear less compacted in the *P_CLB2_*-*MPS2* cell. Red, V5-Csm4; green, Rec8-GFP; blue, DAPI.

We set out to determine the factors that regulate Mps2 binding to the telomere. We found that in the absence of Csm4, Mps2 remained bound to the chromosome ends, indicating that Csm4, which is also localized to the ONM, is not required for Mps2’s association with the telomere (Fig 3C). This finding is consistent with the observation that nuclear localization of Mps2 is independent of Csm4 (Fig 2D). In contrast, removal of Mps3 or Ndj1 abolished Mps2’s binding to the chromosome ends (Fig 3C). Therefore, Mps3 and Ndj1 are required for telomere localization of Mps2. These findings prompted us to hypothesize that Mps2 acts as a linker between Mps3 and Csm4 to mediate t-LINC formation. Indeed, depletion of meiotic Mps2 abolished Csm4’s localization to the telomere (Fig 3C). In contrast, depletion of Mps2 did not alter Mps3’s association with the telomere (Supplemental Fig 2C). Taken together, our findings indicate that the yeast t-LINC is composed of Csm4, Mps2 and Mps3, and that Mps2 links Csm4 and Mps3 together.

### Mps2 regulates telomere bouquet formation and meiotic recombination

At prophase I, the t-LINC mediates telomere bouquet formation and regulates meiotic recombination (Conrad et al., 2008; Kosaka et al., 2008; Koszul et al., 2008; Wanat et al., 2008). To determine the role of Mps2 in meiotic recombination, first, we asked whether Mps2 is required for telomere bouquet formation. We used the telomere-associated protein Rap1 (Conrad et al., 1990), which is tagged with GFP, to serve as a telomere marker. Rap1-GFP formed distinctive foci at the nuclear periphery at prophase I (Fig 4A). In wild-type cells, Rap1 foci were clustered together and often occupied half or less than half of the nuclear periphery, revealing the telomere bouquet configuration at prophase I (Fig 4A and 4B). In contrast, depletion of meiotic Mps2 or removal of Csm4, or both, abolished the telomere bouquet formation (Fig 4A and 4B). Of note, the bouquet configuration appeared to be transient at prophase I. While telomere bouquet took place ubiquitously in individual cells, it was observed in about 25% of the cells in a population 3 hours after the induction of meiosis (Fig 4B). We concluded that like Csm4, Mps2 is required for telomere bouquet formation.

**Figure 4.**
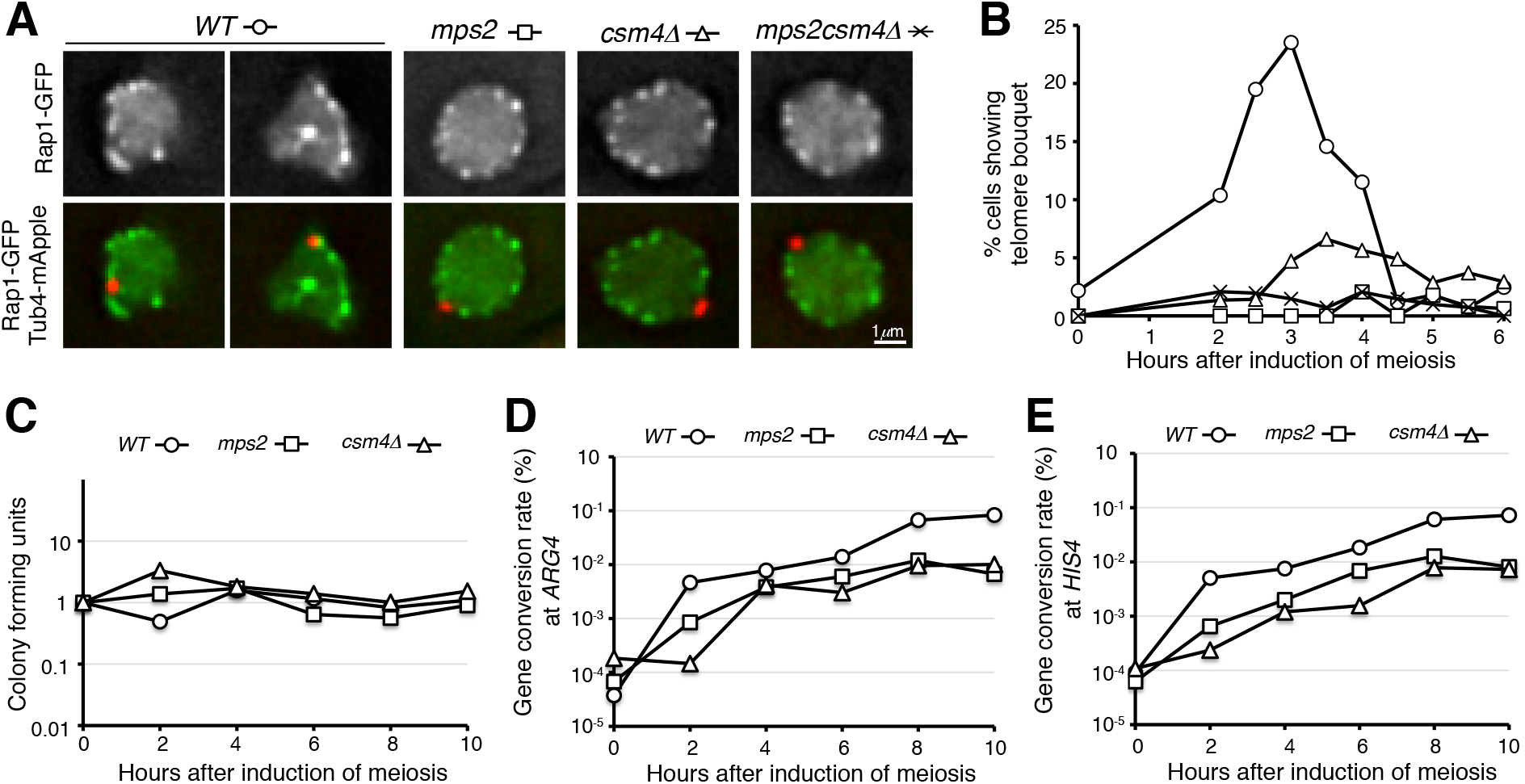
Mps2 regulates telomere bouquet formation. (**A**) Representative images showing Rap1-GFP distribution in *WT*, *P_CLB2_*-*MPS2*, *csm4Δ*, and *P_CLB2_*-*MPS2 csm4Δ* cells at prophase I. Red, Tub4-mApple; green, Rap1-GFP. (**B**) Quantification of telomere bouquet formation in *WT*, *P_CLB2_*-*MPS2*, *csm4Δ*, and *P_CLB2_*-*MPS2 csm4Δ* cells. Three biological replicates were performed; one representative is shown. At least 100 cells were counted at each time point. (**C**-**E**) Gene conversion rate at the *ARG4* and *HIS4* loci in *WT*, *P_CLB2_*-*MPS2*, and *csm4Δ* cells. Yeast cells were induced to undergo synchronous meiosis; aliquots were withdrawn at indicated times. Serially-diluted yeast cells were plated on YPD to determine cell viability (panel C) and on selective dropout medium to determine the rates of gene conversion (panels D and E). Three biological replicates were performed; one representative is shown.

Next, we determined homolog pairing with the TetO/TetR-GFP system that marks chromosome IV at the *LYS4* locus (Supplemental Fig 2D and Jin et al., 2009). In the absence of Mps2, meiotic cells appeared competent to pair at the *LYS4* locus, but displayed a two-hour delay in pairing (Supplemental Fig 2D). This was also the case in *csm4Δ* cells (Supplemental Fig 2D). Finally, we observed that the gene conversion rate at the *ARG4* and *HIS4* loci reduced about 10-fold in both *P_CLB2_*-*MPS2* and *csm4Δ* cells compared to those of the wild type (Fig 4C-E). Together, these findings suggest that Mps2 is required for efficient homolog pairing and meiotic recombination, and further support the notion that Mps2 is a component of the t-LINC complex.

### Reconstitution of t-LINC in vegetative yeast cells

Among the three components of the t-LINC, only Csm4 is specific to meiosis. We therefore hypothesized that by ectopically expressing *CSM4*, the t-LINC would be reconstituted in vegetative yeast cells. We used *P_GAL1_*-*CSM4* to induce Csm4 production in cells grown in galactose medium (Supplemental Fig 3A and 3B). Ectopic expression of *CSM4* caused a slow growth defect (Fig 5A). Crucially, this mutant phenotype was suppressed by the overproduction of Mps2 by way of *P_GAL1_*-*MPS2* (Fig 5A), demonstrating that *CSM4* and *MPS2* genetically interact. In the presence of Csm4, we observed that an ectopic patch of Mps3, but not the SPB marker Tub4, formed in the developing daughter cell during mitosis (Fig 5B-C and Supplemental Fig 3C). Cells with ectopic Csm4 showed a delayed mitotic program (Supplemental Fig 3D), consistent with the slow growth phenotype of *P_GAL1_*-*CSM4* (Fig 5A). Formation of this Mps3 patch corresponded to the precocious extension of the nuclear envelope into the daughter cell before chromosome segregation in mitosis (Supplemental Fig 4A), and crucially, both Mps2 and Mps3 were located at the leading edge of this Csm4-dependent nuclear extension (Supplemental Fig 4B), which is reminiscent of the nuclear protrusions mediated by the t-LINC at meiotic prophase I (Fig 2D and 2E).

**Figure 5.**
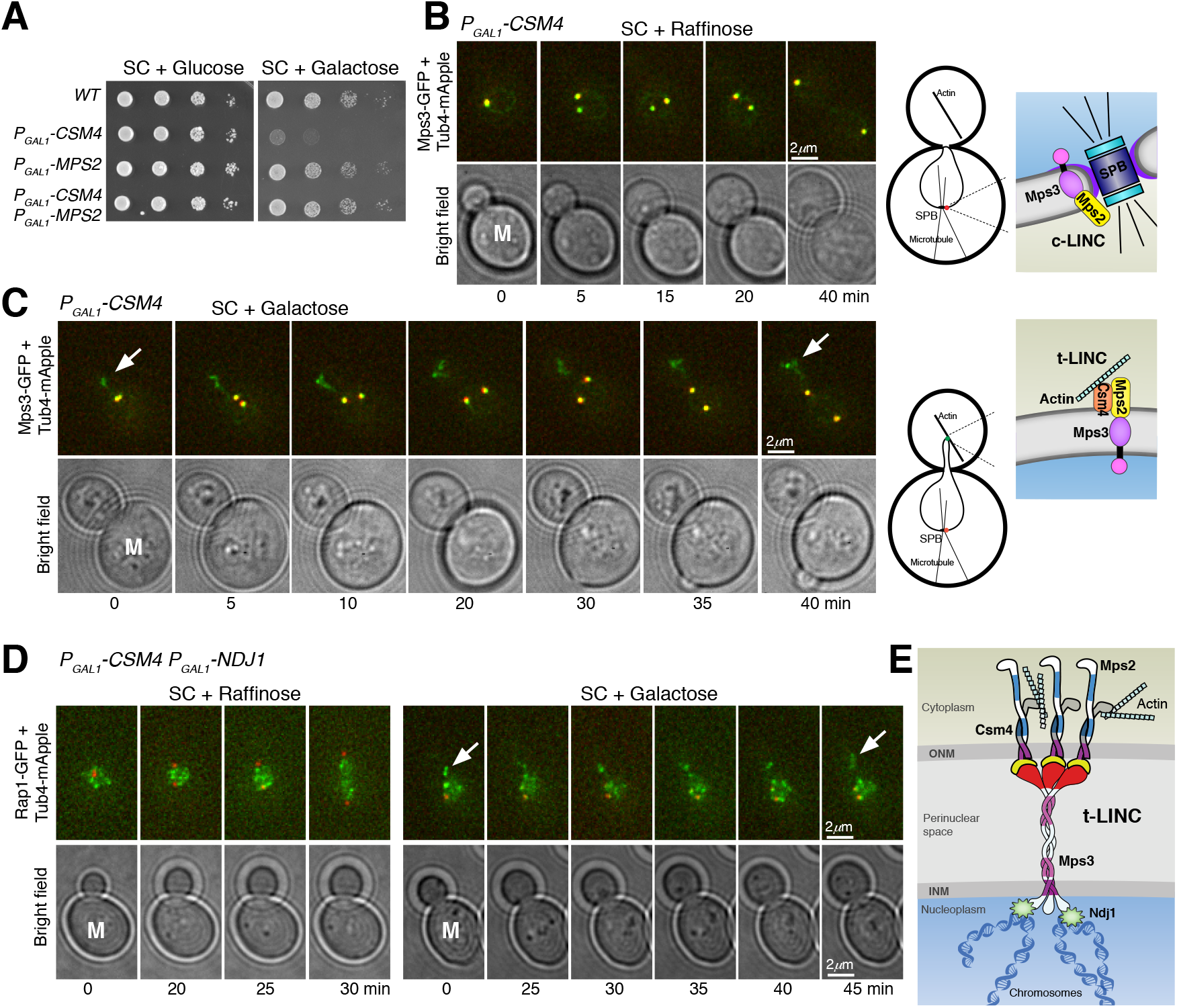
Reconstitution of t-LINC in vegetative yeast cells. (**A**) Genetic interaction between *MPS2* and *CSM4*. Ten-fold diluted yeast cells were spotted onto glucose or galactose medium. Note that ectopic expression of *CSM4* in the galactose medium is toxic to the vegetative yeast cell. SC, synthetic complete. (**B** and **C**) Induction of t-LINC formation in mitosis with ectopic Csm4. Time-lapse fluorescence microscopy showing the localization of Mps3-GFP (green). Tub4-mApple (red) marks the SPB. Arrows pointing to the Mps3 patch formed in the developing daughter cell when Csm4 was produced. Projected images from 12 z sections are shown. Time zero refers to the onset of SPB separation. M, mother cell. Schematic diagrams of c-LINC and t-LINC are shown to the right. (**D**) Reconstituted t-LINC tethers telomeres. Time-lapse fluorescence microscopy was performed as above. Rap1-GFP (green) marks the telomeres; Tub4-mApple (red) marks the SPB. Projected images from 12 z sections are shown. Time zero refers to the onset of SPB separation. Note that in the presence of Csm4, Rap1-GFP formed a patch in the developing daughter cell. (**E**) Model for t-LINC in budding yeast. Three copies of each of Csm4, Mps2 and Mps3 are proposed to form a t-LINC nonamer. INM, inner nuclear membrane; ONM, outer nuclear membrane.

To determine whether the ectopic Mps3 patch depends on Mps2, forming an intact t-LINC complex, we created an *mps2Δ* strain (Supplemental Fig 5). Because *MPS2* is an essential gene, we took advantage of the fact that *pom152Δ* suppresses the lethal phenotype of *mps2Δ*, presumably bypassing the need of Mps2 in SPB duplication (Supplemental Fig 5A and Katta et al., 2015). In *mps2Δ pom152Δ* double mutant cells, we never observed the ectopic Mps3 patch in the daughter cell with or without the presence of Csm4 (Supplemental Fig 5B). Together, these findings demonstrate that both Csm4 and Mps2 are required for the formation of the ectopic Mps3 patch, therefore the intact t-LINC complex, in the daughter cell during mitosis.

Because t-LINC mediates actin-based motility in meiosis (Koszul et al., 2008), we reasoned that formation of the ectopic Mps3 patch and thereby the t-LINC depends on actin polymerization, which is highly active in the budding daughter cell (Bi and Park, 2012). To test this hypothesis, we treated yeast cells with the actin polymerization inhibitor Latrunculin B (Lat B) (Supplemental Fig 3E). Mps3 patches disappeared 15 minutes after the addition of Lat B to the yeast medium (Supplemental Fig 3E), suggesting that formation of the ectopic t-LINC in the daughter cell depends on actin polymerization. Together, these findings demonstrate that t-LINC can be reconstituted in vegetative yeast cells simply by ectopic production of Csm4 and that Mps2 acts as the linker that connects Csm4 to Mps3, confirming the heterotrimeric nature of the t-LINC in budding yeast.

### Reconstituted t-LINC can tether telomeres

To determine whether reconstituted t-LINC is capable of tethering telomeres, we induced the production of Ndj1 together with Csm4 in vegetative yeast cells (Fig 5D). We have shown previously that Ndj1, when ectopically produced in mitosis, binds to Mps3 (Li et al., 2015). In the presence of Ndj1, we predicted that the t-LINC tethers telomeres to the nuclear envelope. In wild-type cells, telomeres, marked by Rap1-GFP, trailed the separating SPBs during mitosis (Fig 5D). In contrast, in the presence of both Csm4 and Ndj1, Rap1-GFP entered the daughter cell precociously, forming an ectopic patch just like the Mps3 patch, well before SPB separation (Fig 5D). Therefore, ectopically reconstituted t-LINC is functional in tethering telomeres.

In conclusion, we have demonstrated the heterotrimeric composition of the budding yeast t-LINC; specifically, the KASH-like protein Mps2 bridges Csm4 and Mps3 (Fig 5E). Four lines of evidence support the idea of a heterotrimeric nature of the yeast t-LINC. First, Mps2 is a major binding partner of Csm4 and colocalizes with Csm4 at the telomere; second, Mps2 is required for Csm4’s association with the telomere, but not for Mps3; third, Mps2 is essential for telomere bouquet formation, a major activity of t-LINC; and finally, Mps2 is required for functional reconstitution of t-LINC in vegetative yeast cells by linking Csm4 and Mps3 together. We note that a recent work also suggests that Mps2 acts as a member of the t-LINC (Lee et al., 2020). In budding yeast, the combined action of Mps2 and Csm4 is needed to carry out the function of the canonical KASH protein at the t-LINC. On the basis of the current understanding of the oligomerization state of SUN and KASH proteins (Sosa et al., 2012; Wang et al., 2012; Jahed et al., 2018), we propose that the yeast t-LINC is a nonamer (Fig 5E). In Arabidopsis, WIP and WIT proteins are located at the ONM and appear to interact with each other to function as a KASH protein (Meier, 2016), which is analogous to Mps2 and Csm4. In metazoans, numerous KASH variants also exist. Our work suggests that variant LINC complexes could be prevalent and provides insight into LINC assembly and its evolution in eukaryotes.

## Materials and methods

### Yeast strains and plasmids used in this study

Yeast strains and plasmids used in this study are listed in Tables S1 and S2. Strains for meiotic experiments are isogenic to the SK1 genetic background and strains for mitotic experiments are from the S288C background. To generate proteins that are tagged at their N-termini, alleles of *TAP-MPS2*, *V5-CSM4*, *TAP-CSM4*, *V5-MPS2*, *TAP-MPS3*, *GFP-MPS2* and *GFP-CSM4* were created by homologous recombination-based gene replacement as we have described previously (Koch et al., 2019). Briefly, the corresponding plasmids (Table S2) were linearized by restriction digestion and integrated at the endogenous locus of each respective allele by yeast transformation. To remove the untagged gene, *URA3* positive colonies were then counter-selected on a 5’-FOA plate; these tagged alleles therefore served as the only functional copy in the yeast genome. All of these alleles were verified by DNA sequencing before use.

To generate C-terminal tagged alleles, a PCR-based yeast transformation method (Longtine et al., 1998) was used to generate *CSM4-GFP*, *MPS2-RFP*, and *MPS2-3HA*. Positive transformants were confirmed by colony PCR. A comparable PCR-based method was used to replace the *CSM4*, *MPS2* and *POM152* open reading frames with either a KanMX4 or Hygromycin-B cassette to generate gene deletions. Correct transformations were further confirmed by colony-based diagnostic PCR. Using a similar PCR-based method, *P_CLB2_*-*MPS2* was generated by replacing the endogenous promoter with the mitosis-specific promoter from *CLB2* (Lee and Amon, 2003). Primers used in this study are included in Table S3. The following alleles have been reported previously: *MPS3-V5*, *ndt80Δ*, *MPS3-mApple*, *TUB4-mApple*, *NDJ1-V5*, *HTA1-mApple*, *REC8-GFP*, *P_CLB2_*-*MPS3*, *ndj1Δ*, *RAP1-GFP* and *MPS3-GFP* (Li et al., 2014; Li et al., 2015).

To ectopically express *CSM4* in vegetative yeast cells we constructed *P_GAL1_*-*V5-CSM4* (pHG317) to express the full length *CSM4* under the control of the *GAL1* promoter. We linearized plasmid pHG317 with PstI and integrated it at the endogenous *CSM4* locus by yeast transformation. A similar approach was used to overexpress *MPS2* in vegetative yeast cells by constructing *P_GAL1_*-*GFP-MPS2* (pHG527). Plasmid pHG527 was linearized with StuI and integrated at the endogenous *MPS2* locus by yeast transformation. Note that the endogenous *MPS2* remains intact and functional. The plasmid pHG335 (*P_GAL1_*-*V5-NDJ1*) has been described previously (Li et al., 2015). Leucine positive colonies were confirmed by colony-based diagnostic PCR.

### Yeast culture method and cell viability assay

For meiotic experiments, yeast cells were grown in YPD (1% yeast extract, 2% peptone, and 2% dextrose) at 30OC. These YPD cultures were diluted with YPA (1% yeast extract, 2% peptone, and 2% potassium acetate) to reach OD (optical density, λ = 600nm) of 0.2 and incubated at 30OC for approximately 14 hours to reach a final OD of ~1.6-1.8. Yeast cells were then washed by water and resuspended in 2% potassium acetate to induce synchronous meiosis as described previously (Koch et al., 2019). Yeast samples were withdrawn at the indicated times for fluorescence microscopy and/or protein extraction.

To synchronize cycling cells, S288C yeast cells were grown in SC (synthetic complete) medium with 2% raffinose to OD of 0.5 and arrested at G1 phase by the addition of 10 μg/ml (final concentration) of alpha factor. Raffinose cultures were separated into 2 flasks after the addition of alpha factor; one served as the control and the other one received galactose (2% final concentration) 30 minutes before alpha factor removal. The addition of galactose induced the expression of *GAL*-regulated genes prior to the release of yeast cells from G1 arrest. To remove alpha factor, cells were washed twice with water and once with SC raffinose or galactose, and then resuspended in the respective medium. Samples were withdrawn at the indicated times for fluorescence microscopy and/or protein extraction.

To determine cell growth, yeast cells were grown overnight to reach saturation in YPD liquid medium, 10-fold diluted, spotted onto SC plates containing either 2% glucose or 2% galactose and then incubated at 30°C for about two days.

### Protein affinity purification and mass spectrometry

Protein affinity purification was performed as we have reported previously (Li et al., 2015). In brief, 2 liters of yeast cells were induced into synchronous meiosis for 6 hours. Yeast cells were harvested and ground into powder in the presence of liquid nitrogen. The yeast powder was then stored at −80°C before use. For affinity purification, yeast powder was thawed in the extraction buffer. The lysate was then incubated with epoxy-activated M-270 Dynabeads (Thermo Fisher Scientific, Cat#14305D), which were cross-linked with rabbit IgG (Sigma-Aldrich, Cat#I5006). The final product was eluted from the beads and dried for further study. Purified protein samples were digested by trypsin. The proteomics work was carried out by the Translational Science Laboratory, Florida State University College of Medicine. An externally calibrated Thermo LTQ Orbitrap Velos mass spectrometer was used for mass spectrometry per the method described previously (Li et al., 2015).

### Protein extraction and western blotting

For meiotic yeast cells, proteins were extracted with the trichloroacetic acid (TCA) method as described previously (Jin et al., 2009). In brief, 3 to 5 mL of yeast cells were collected, resuspended in 2.5% ice cold TCA, and incubated at 4°C for 10 minutes. Cell pellets were stored at −80 °C before use, and proteins were extracted in the RIPA buffer by bead beating with a mini bead-beater homogenizer for 90 seconds at 4°C prior to standard SDS-PAGE and western blotting.

For mitotic experiments, yeast aliquots were withdrawn at the indicated times for protein extraction by precipitation in the presence of 20 mM NaOH and standard SDS-PAGE and western blotting protocols were followed (Koch et al., 2019).

Proteins tagged with HA (Mps2-3HA and 3HA-Mps2) were detected by an anti-HA mouse monoclonal antibody (1:1,000 dilution, 12CA5; Sigma). Similarly, V5-tagged proteins (V5-Csm4, Mps2-V5, and Mps3-V5) were detected by an anti-V5 mouse monoclonal antibody (1:1,000 dilution, Proteintech, cat#66007-1-Ig), and TAP-tagged proteins (TAP-Csm4, TAP-Mps2, and TAP-Mps3) were detected by an anti-TAP rabbit antibody (1:10,000, Thermo Fisher Scientific, cat#CAB1001). The level of Pgk1 was detected by a Pgk1 antibody (1:10,000, Thermo Fisher Scientific, cat#PA5-28612) and was used as a loading control. Horseradish peroxidase-conjugated secondary antibodies, goat anti-mouse and goat anti-rabbit (Bio-Rad, cat#1706516 and 1705046), were used to probe the proteins of interest by an ECL kit (Bio-Rad, cat#1705060). Two ECL-based western blot detection methods were utilized, X-ray film (Figs 1-5) and the ChemiDoc MP Imaging System (Bio-Rad, cat#17001402) (Supplemental Fig 3).

### Live-cell fluorescence microscopy

Live-cell fluorescence microscopy was conducted on a DeltaVision imaging system (GE Healthcare Life Sciences) with a 63× objective lens (NA=1.40) on an inverted microscope (IX-71; Olympus). Microscopic images were acquired with a CoolSNAP HQ2 CCD camera (Photometrics). Prior to microscopy, yeast cells were prepared as described previously (Li et al., 2015). Briefly, yeast cells were prepared on a concave microscope slide (approximately 0.8 mm deep) filled with an agarose pad with 2% potassium acetate. The concave slide was then sealed with a cover slip and scoped for the desired time duration. The microscope stage was enclosed in an environmental chamber set at 30°C. For time-lapse microscopy, optical sections were set at 0.5 μm thickness with 12 z sections. Ultra-high signal-to-background coated custom filter sets were used. For GFP, the excitation spectrum was at 470/40 nm, emission spectrum at 525/50 nm; for RFP, excitation was at 572/35 nm, and emission at 632/60 nm. To minimize photo toxicity to the cells and photo bleaching to fluorophores, we used neutral density filters to limit excitation light to 32% or less of the normal equipment output for time-lapse microscopy. Images were deconvolved with SoftWoRx (GE Healthcare Life Sciences); projected images or single optical sections were used for display.

To determine meiotic cell progression, aliquots of yeast cells were collected at indicated times and prepared for fluorescence microscopy. Tub4-mApple serves as an SPB marker. At least 100 cells were counted at each time point to determine the rate of SPB separation.

In experiments testing the dependence of actin filaments, Latrunculin B (final concentration of 100 μM) was added to the cell culture prior to microscopy. The same volume of DMSO was added in the control group.

### Nuclear spread and immunofluorescence

Surface nuclear spreads were performed as described previously (Jin et al., 2009). In brief, yeast cells enriched at prophase I (~5 hours after induction of meiosis) were spheroplasted by lyticase treatment. Spheroplasts were then fixed and poured onto a glass slide. The slide was then rinsed with PhotoFlo 200 and air dried, followed by PBS buffer with 3% BSA to block for 2 hours at room temperature. Anti-V5 antibody (R960-25; Thermo Fisher Scientific) was used to detect V5-Csm4 and Ndj1-V5; anti-HA antibody (12CA5; Roche/Sigma) was used to detect Mps2-3HA. Rec8-GFP was detected by an anti-GFP mouse monoclonal antibody (ab209, Abcam). Secondary antibodies (FITC-conjugated goat anti–rabbit, rhodamine-conjugated goat anti–mouse, and Cy3-conjugated goat anti–rat; Jackson ImmunoResearch Laboratories) were used at a dilution of 1:500. Mounting medium with DAPI was added before microscopy. Images were acquired with an epifluorescence microscope (Axio Imager M1, Zeiss) with 100× objective lens (NA=1.40) at room temperature.

### Gene conversion assay

Yeast cells were induced to undergo synchronous meiosis and aliquots were withdrawn at the indicated times. Serially diluted yeast cells were plated on YPD plates to determine cell viability, and on SC Arginine-dropout and SC Histidine-dropout plates to determine gene conversion rate at the *ARG4* and *HIS4* loci. The rate of gene conversion was calculated by the ratio of the colony-forming units on SC dropout plates over those of on YPD plates.

## Acknowledgement

We thank Hank Bass and Yanchang Wang for discussions. Elizabeth Staley provided technical support. Jen Kennedy and Charles Badland assisted with text editing and graphic design. This work was supported by the National Institute of General Medicine (GM117102) and the National Science Foundation (MCB1951313).

## Supplemental information

Supplemental table 1. Yeast strains used in this study

Supplemental table 2. Plasmids used in this study

Supplemental table 3. Primers used in this study

## Supplemental figure legends

**Supplemental Figure 1.**
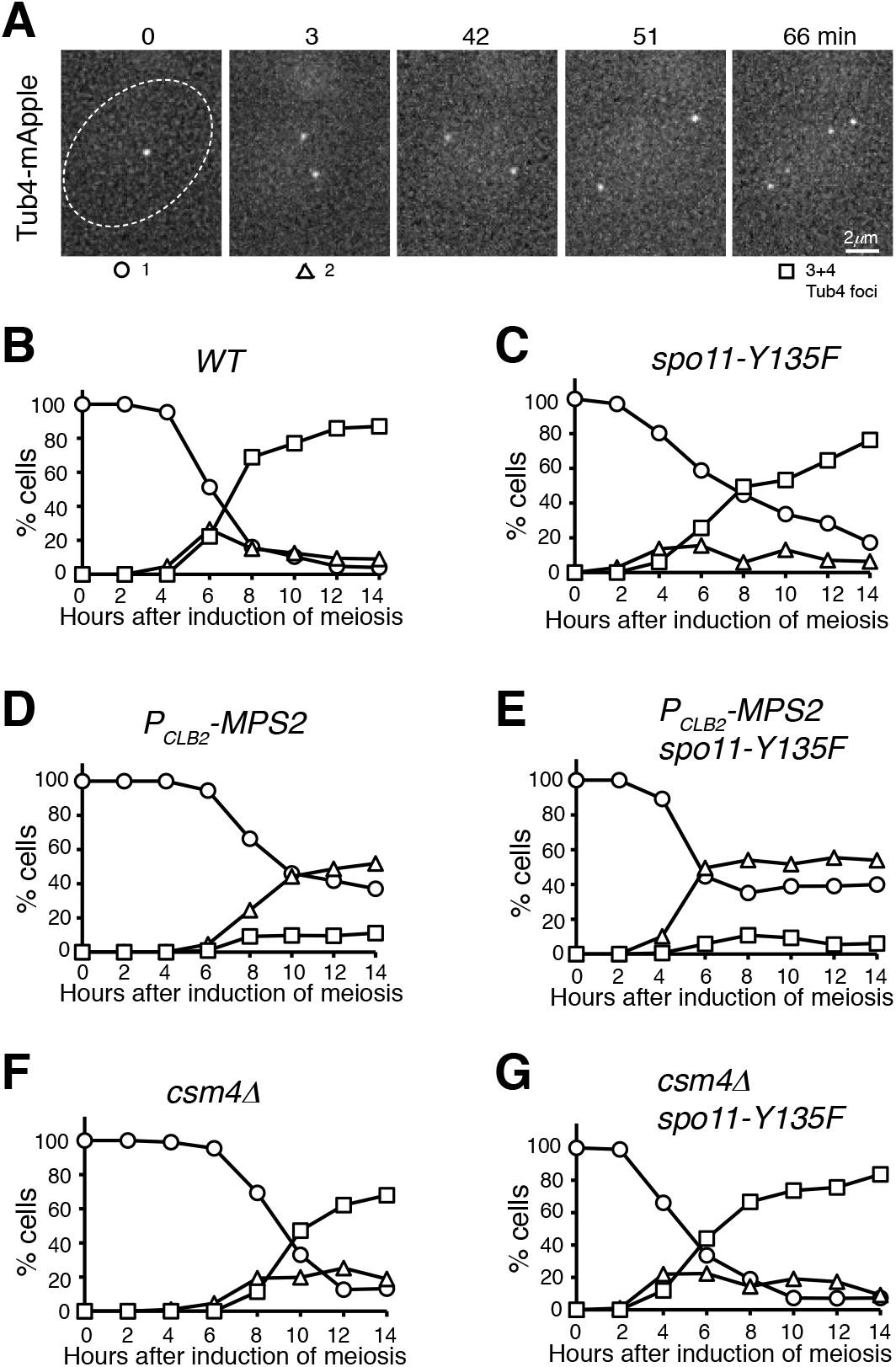
SPB separation in budding yeast meiosis. (**A**) Time-lapse fluorescence microscopy showing a representative cell undergoing meiosis. Tub4-mApple marks the SPB. Time zero refers to the point of SPB separation in meiosis I. Time in minutes is shown at the top. (**B**-**G**) Quantification of SPB separation in *WT* (**B**), *spo11-Y135F* (**C**), *P_CLB2_*-*MPS2* (**D**), *P_CLB2_*-*MPS2 spo11-Y135F* (**E**), *csm4Δ* (**F**) and *csm4Δ spo11-Y135F* (**G**) cells during meiosis. At least 100 cells were counted at each time point. Three biological replicates were performed; one representative is shown.

**Supplemental Figure 2.**
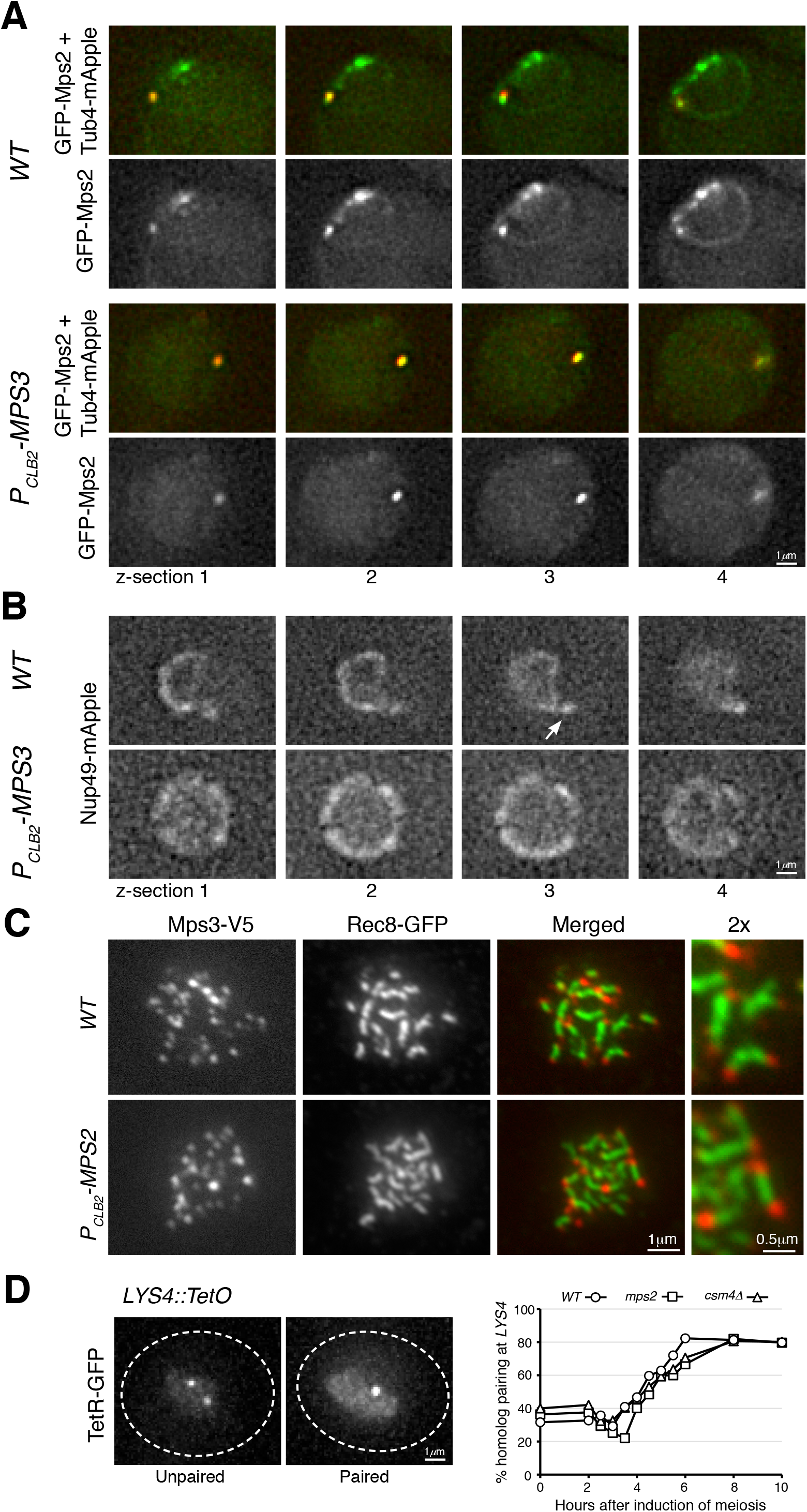
Mps2 localization and homolog pairing in meiosis. (**A**) Live-cell fluorescence microscopy showing GFP-Mps2 (green) localization at prophase I. Note that GFP-Mps2 is clustered around the nuclear periphery in the wild-type (WT) cell but is only visible at the SPB in the *P_CLB2_*-*MPS3* cell. Four continuous optical sections are shown. Tub4-mApple (red) marks the SPB. (**B**) Live-cell fluorescence microscopy showing Nup49-mApple localization at prophase I. The arrow points to the nuclear protrusion in the wild-type cell. Four continuous optical sections are shown. (**C**) Meiotic Mps3 binds to telomeres. Surface nuclear spreads were prepared as in Fig 3. Note that Mps3 remains bound to chromosome ends in the Mps2-depleted cell. Rec8 marks the chromosome axis. Red, Mps3-V5; green, Rec8-GFP. (**D**) Quantification of homolog pairing at the *LYS4* locus. Cells were induced to undergo synchronous meiosis; aliquots were withdrawn at indicated times. TetR-GFP forms a nuclear focus when chromosome IV homologs are paired at *LYS4*. At least 100 cells were counted at each time point. Three biological replicates were performed; one representative is shown.

**Supplemental Figure 3.**
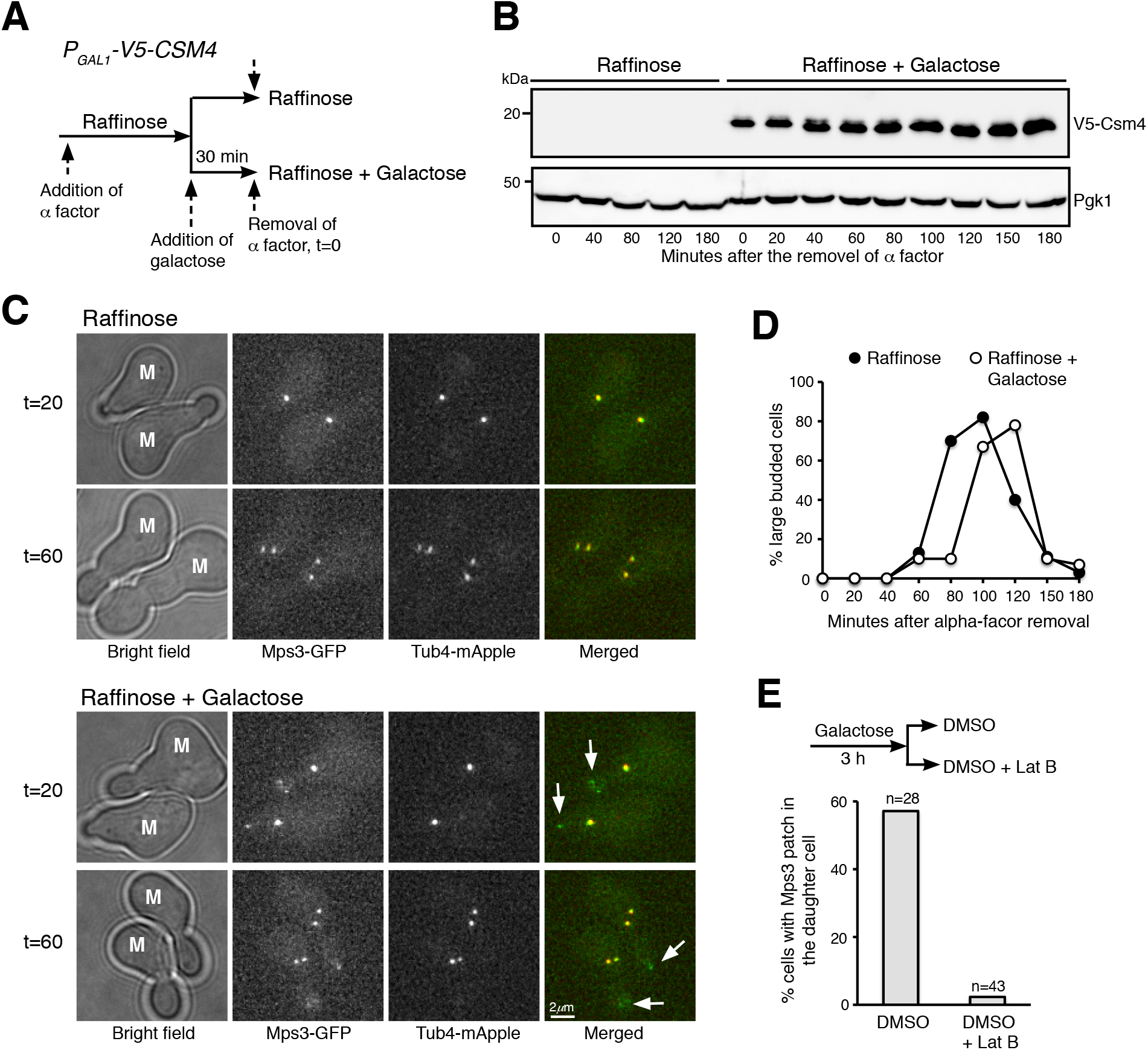
Ectopic production of Csm4 reconstitutes t-LINC in mitosis. (**A**) Schematic diagram showing the experimental procedure. (**B**) Western blotting showing induced production of Csm4 in the galactose medium. V5-Csm4 was probed by an anti-V5 antibody. The level of Pgk1 serves as a loading control. (**C**) Formation of the Mps3 patch in the daughter cell in the presence of Csm4. Tub4-mApple (red) marks the SPB. Projected images of 12 z-sections are shown. Arrows point to the Mps3-GFP (green) patch in daughter cells. M, mother cell. (**D**) Quantification of budding index. Cell aliquots were withdrawn at indicated times and budding morphology was determined by phase contrast microscopy. More than 200 cells were counted at each time point in both raffinose and galactose treatments. (**E**) Impact of actin polymerization on Mps3 patch formation. Schematic diagram at the top shows the experimental procedure. Fluorescence microscopy was performed 15 minutes after the treatments to visualize Mps3-GFP patch formation as in panel C. Lat B, Latrunculin B.

**Supplemental Figure 4.**
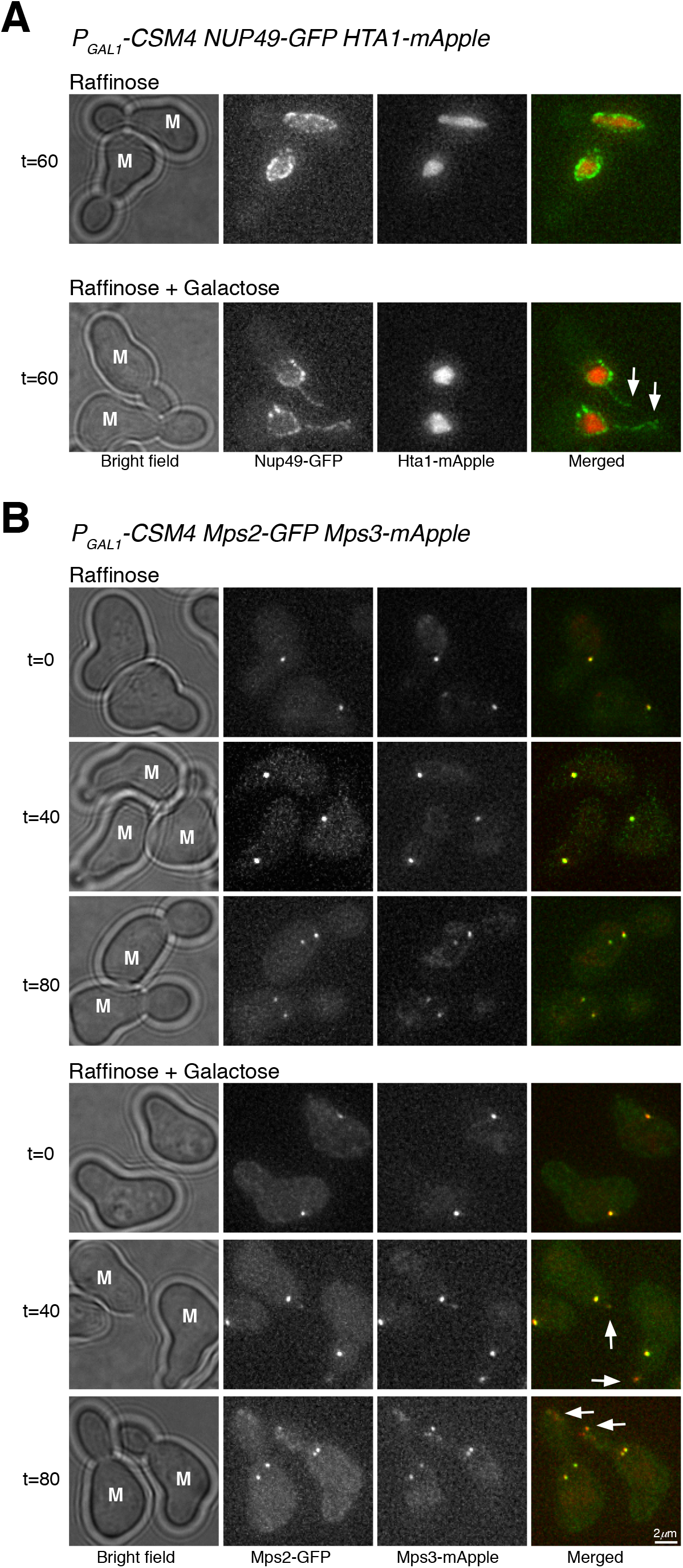
Extension of the nuclear envelope and colocalizaiton of Mps2 and Mps3 at the reconstituted t-LINC. (**A**) Ectopic expression of *CSM4* leads to precocious extension of the nuclear envelope into the daughter cell during mitosis. Induction of *P_GAL1_*-*CSM4* was performed as in Supplemental Fig 3A. Cells were collected at indicated times for live-cell fluorescence microscopy. Nup49, a nuclear pore complex component, marks the nuclear envelope. Hta1 is the yeast histone H2A. Note that in the presence of Csm4, Nup49 and therefore its associated nuclear envelope forms an extended line into the daughter cell (arrows). Red, Hta1-mApple; green, Nup49-GFP. (**B**) Colocalization of Mps2 and Mps3 at the reconstituted t-LINC. Note that in the presence of Csm4, Mps2 and Mps3 colocalize to the leading edge of the nuclear extension in the daughter cell (Arrows). In the mother cell (M), Mps2 and Mps3 colocalize at the SPB. Red, Mps3-mApple; green, Mps2-GFP.

**Supplemental Figure 5.**
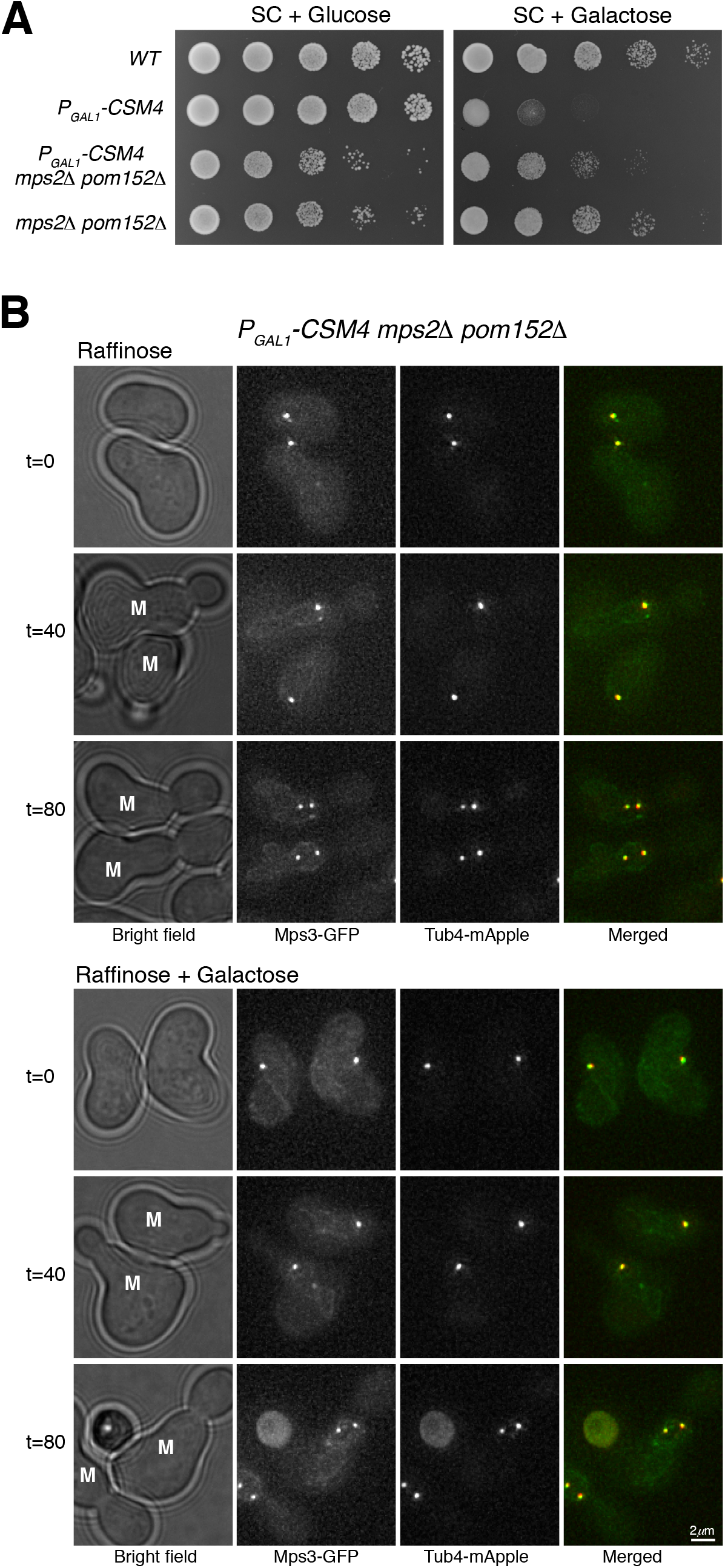
Mps2 is required for bridging Csm4 and Mps3 to reconstitute the t-LINC. (**A**) Cell growth assay. Ten-fold diluted yeast cells were spotted onto raffinose or galactose medium. Note that in the absence of Mps2 and Pom152, Csm4 is no longer toxic in vegetative yeast cells. (**B**) Mps2 is required for forming the Mps3 patch in daughter cells. A similar experimental procedure was carried out as shown in Supplemental Fig 3A. Live-cell fluorescence microscopy was performed as in Supplemental Fig 3C. Note that Mps3-GFP no longer forms a patch in the daughter cell when Mps2 is absent. Red, Tub4-mApple; green, Mps3-GFP. M, mother cell.

---

**Table S1.**
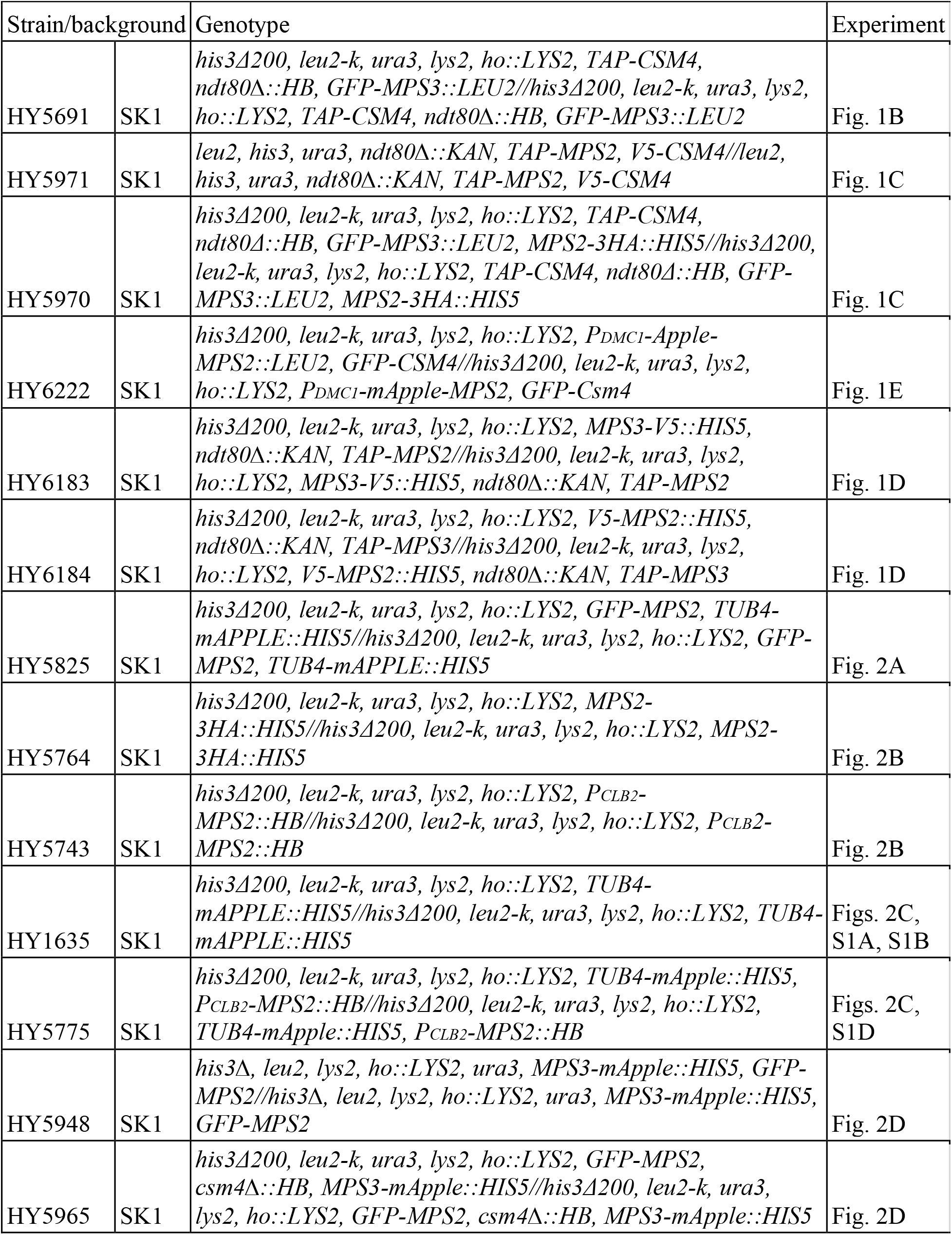
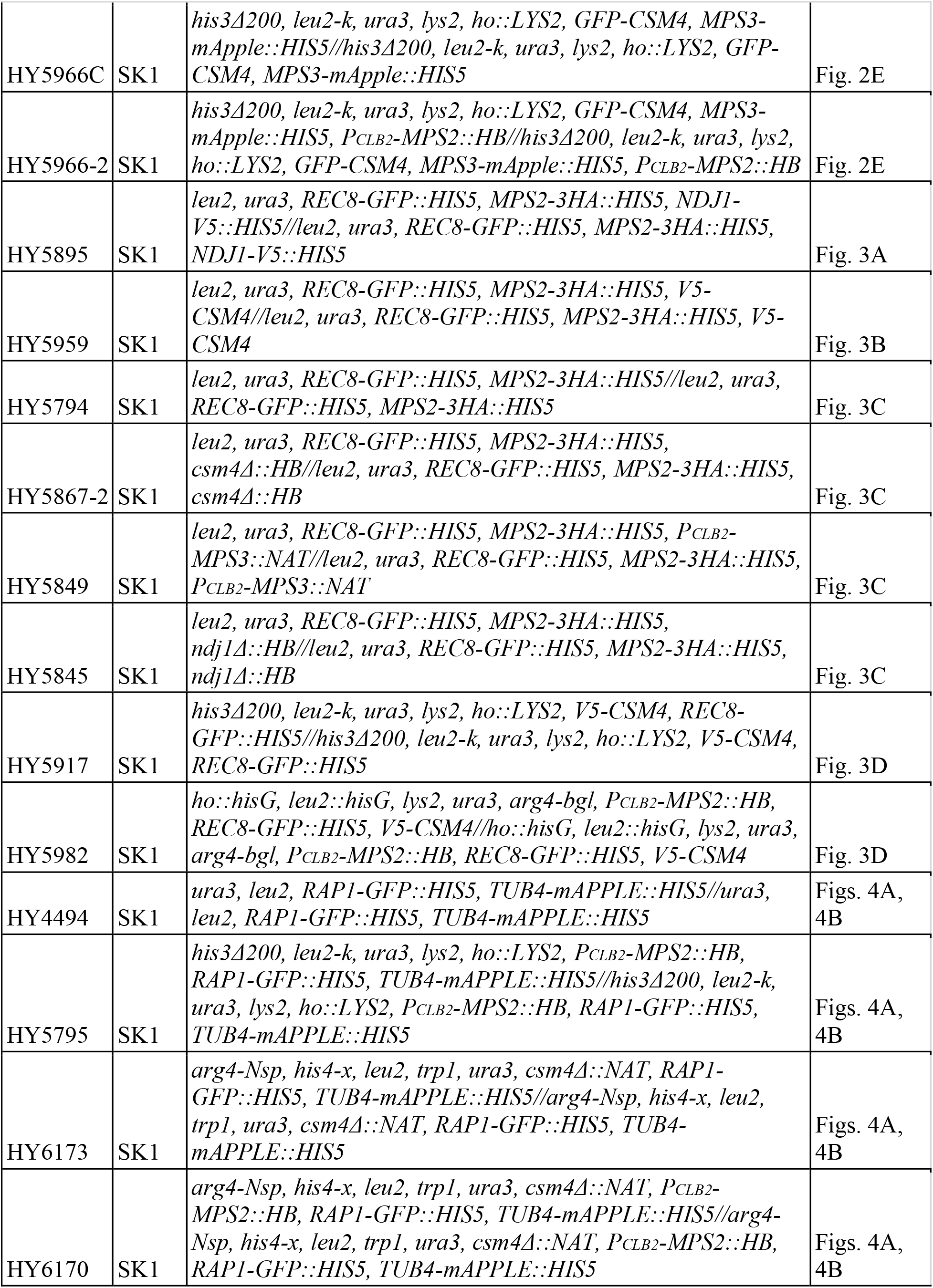
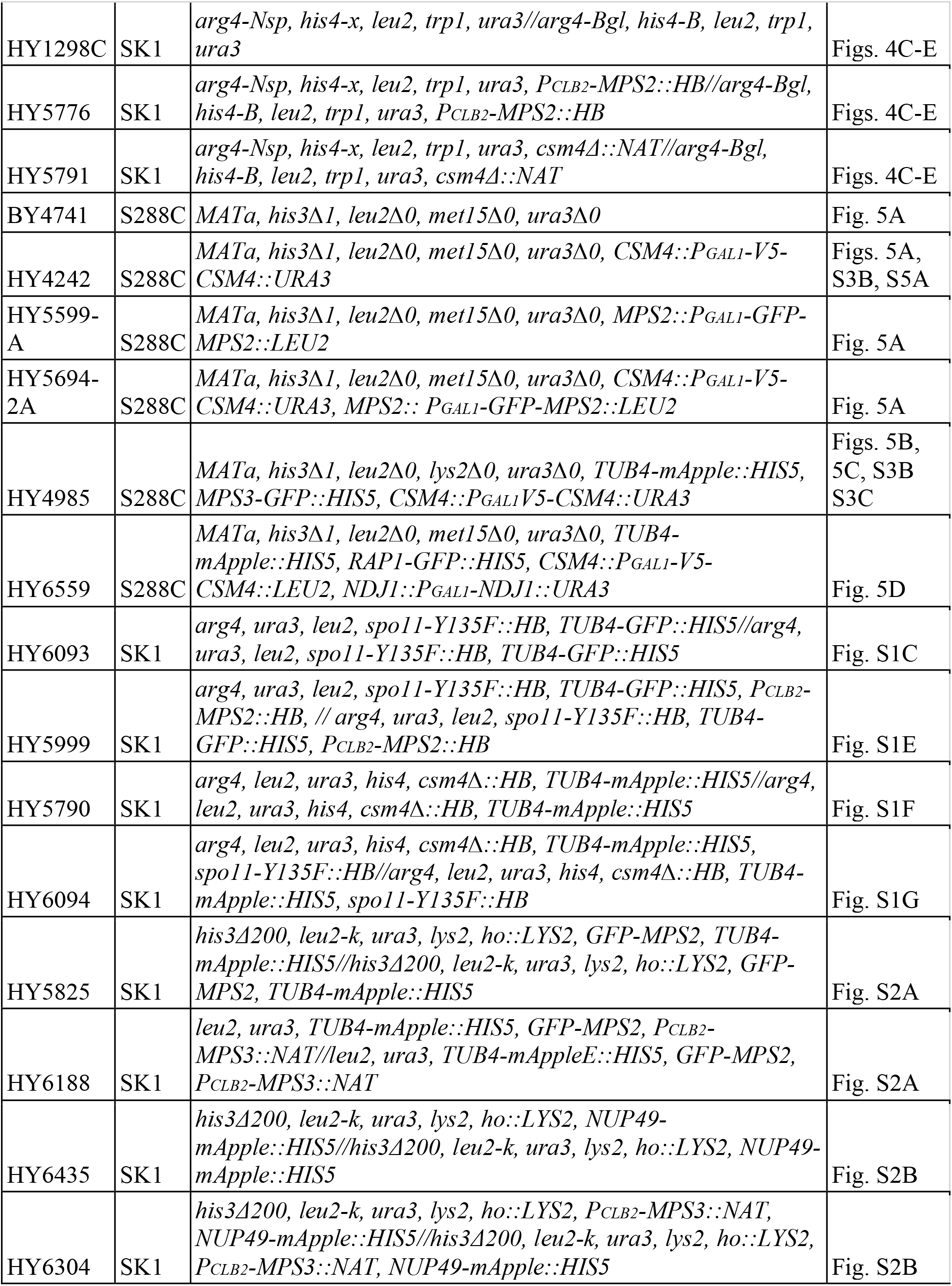
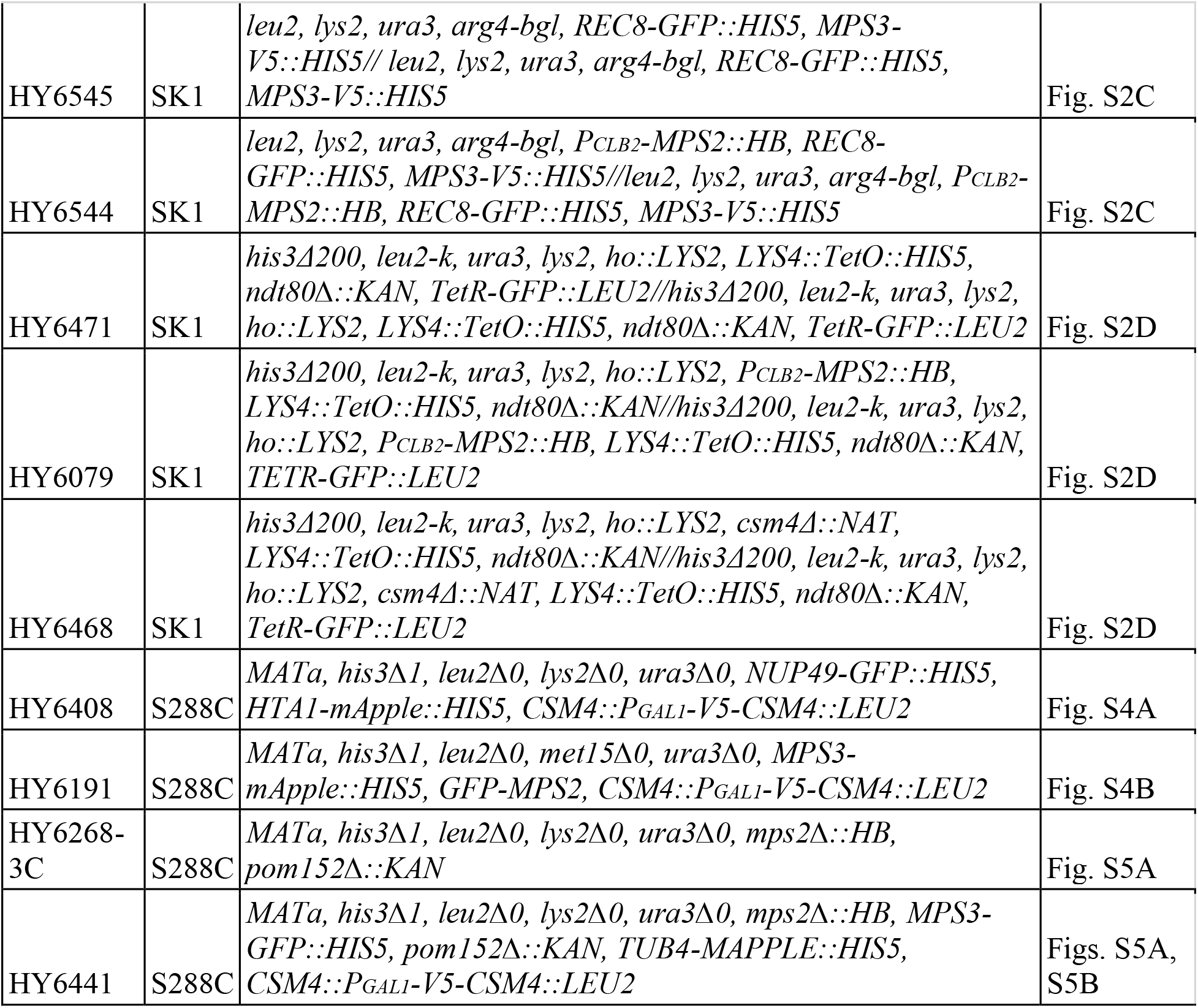
Yeast strains used in this study.

**Table S2.** Plasmids used in this study.

| Plasmid name | Gene construct | Selective marker |
| --- | --- | --- |
| pHG521 | *P<sub>CSM4</sub>-TAP-CSM4* | *URA3* |
| pHG563 | *P<sub>MPS2</sub>-TAP-MPS2* | *URA3* |
| pHG564 | *P<sub>CSM4</sub>-V5-CSM4* | *URA3* |
| pHG581 | *P<sub>MPS2</sub>-V5-MPS2* | *URA3* |
| pHG562 | *P<sub>MPS2</sub>-GFP-MPS2* | *URA3* |
| pHG476 | *P<sub>CSM4</sub>-GFP-CSM4* | *LEU2* |
| pK3524 | *TetR-GFP* | *LEU2* |
| pHG334 | *P<sub>GAL1</sub>-V5-CSM4* | *URA3* |
| pHG527 | *P<sub>GAL1</sub>-GFP-MPS2* | *LEU2* |
| pHG315 | *P<sub>GAL1</sub>-V5-NDJ1* | *LEU2* |
| pHG317 | *P<sub>GAL1</sub>-V5-CSM4* | *LEU2* |

**Table S3.**
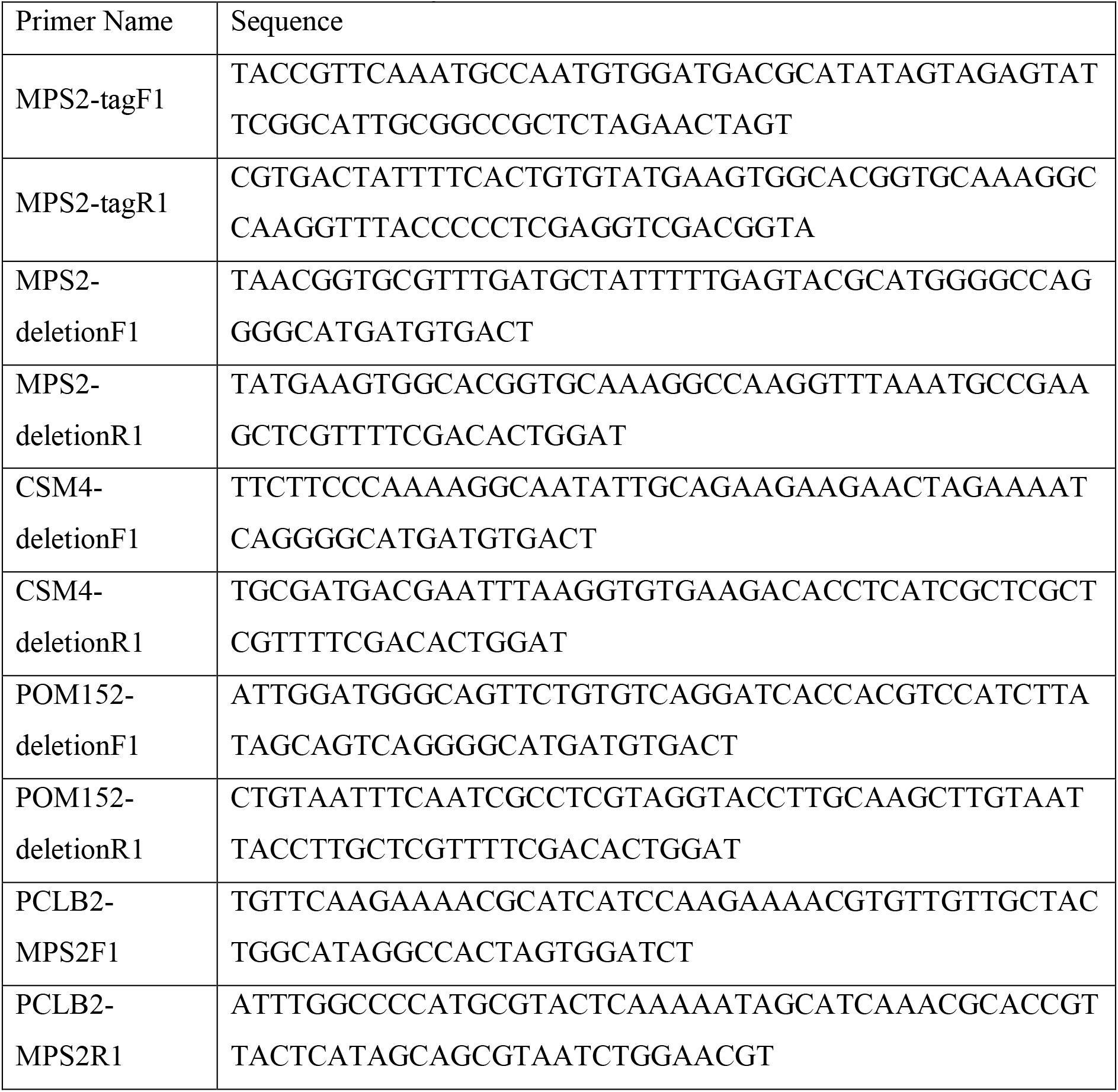
Primers used in this study.

